# Rediscovery of the Enigmatic *Paroxygraphis* in Xizang, China: Phylogenetic Evidence for its Reclassification Within *Oxygraphis* (Ranunculaceae)

**DOI:** 10.1101/2025.07.16.665131

**Authors:** Wen-He Li, Wan-Ting Chen, Bo-Wen Liu, Jia-Min Xiao, Huan-Yu Wu, Jian He, Lei Xie

**Affiliations:** School of Ecology and Nature Conservation, Beijing Forestry University, Beijing 100083, China

**Author notes:** These authors contributed equally to this work (Wen-He Li, Wan-Ting Chen). **Authors for correspondence:** Jian He; Lei Xie.

**Keywords:** molecular phylogeny, morphology, new record, Ranunculeae, taxonomy

## Abstract

The enigmatic monotypic genus *Paroxygraphis* (Ranunculaceae, trib. Ranunculeae), represented solely by *P. sikkimensis*, has long posed taxonomic uncertainties due to its rarity. Through field rediscovery in southern Xizang province, China, and comprehensive molecular phylogenetic analyses combining nrITS and four plastid regions, we resolve its phylogenetic position within the *Oxygraphis* clade, with an estimated divergence time of about 3.73 Ma (95% HPD: 1.98–5.77 Ma) from its sister clade. Morphological and ecological characteristics, including small herb with a rosette of basal simple leaves, longitudinally ribbed achenes, and its alpine adaptations, further support this relationship. Furthermore, the results showed that persistent sepals and unisexual flowers in trib. Ranunculeae reflects multiple independent evolutionary origins. We propose the new combination *Oxygraphis sikkimensis* and highlight its critically endangered status due to habitat vulnerability. Lectotypification of *P. sikkimensis* is also made. This study provides new data for the floristic research of China and resolves the long-standing taxonomic controversy of this problematic genus within trib. Ranunculeae.

## 1 Introduction

The buttercup family, Ranunculaceae, occupying a critical phylogenetic position within the eudicot clade (Chase et al. 2016), has long been a focus of botanical research. Bibliometric analyses indicate that Ranunculaceae is one of the most extensively studied basal eudicot families, with over 55,000 publications indexed in Google Scholar and over 18,000 publications indexed in Web of Science since 1950.

This sustained research interest arises from its evolutionary significance as an early-diverging lineage, its nearly cosmopolitan distribution across diverse ecosystems (Wang et al. 2016), and its economic importance due to ornamental and medicinal applications (Ro et al. 1997). Nevertheless, several genera within Ranunculaceae remain insufficiently understood and continue to pose taxonomic challenges.

The monotypic genus *Paroxygraphis* W. W. Smith (Ranunculaceae, trib. Ranunculeae) was first described in 1913 (Smith 1913), representing a unique taxonomic entity within the buttercup family (Tamura 1995). The sole species of the genus, *P. sikkimensis* W. W. Smith, is a small and inconspicuous herbaceous plant characterized by fleshy fibrous roots, simple leaves arranged in a basal rosette ( with long petioles and notably small blades), and most diagnostically dioecious reproduction, a rare trait among its predominantly hermaphroditic relatives. This genus is distributed in the eastern Himalayas at elevations of 3000–4000 m. Due to its diminutive size and extremely restricted distribution, *P. sikkimensis* has been poorly documented and understood since its initial description.

Taxonomic treatments of Ranunculaceae, notably Tamura (1995) and Emadzade et al. (2010), consistently recognize *Paroxygraphis* as a distinct genus within trib. Ranunculeae. This tribe represents one of the most diversed lineages within the Ranunculaceae family, comprising 18 recognized genera (Emadzade et al. 2010) and roughly 650 currently known species (Tamura 1993, 1995). The tribe Ranunculeae has unitegmic ascending ovules (except *Myosurus* L. which has pendent ovules; Tamura 1995) and petals with at least one nectary gland near the base. Within Ranunculeae, *Ranunculus* L. dominates with over 600 species (Hörandl and Emadzade 2011), while the remaining 17 genera are smaller, including several monotypic ones largely confined to cold temperate regions of the Americas and Europe (Tamura 1995).

Based on gross morphology, distribution patterns, and ecological preferences, *P. sikkimensis* strongly suggest a close relationship with the genus *Oxygraphis* Bunge, a small Ranunculeae genus of four or five species (Figure 1) inhabiting Asian alpine to subarctic zones (Tamura 1995; Wang et al. 2001; Rai and Rawat 2015). All species of *Oxygraphis* are acaulescent perennial herbs with fleshy fibrous roots, and their leaves are exclusively basal, simple, and petiolate. The flowers are bisexual, characterized by multiple petals (typically 5–15) and numerous pistils (15–40). The fruits are longitudinally ribbed achenes (Tamura 1995; Wang et al. 2001). Four *Oxygraphis* species occur in China (Wang et al. 2001), and like *P. sikkimensis*, most (three or four species) retain persistent sepals post-anthesis—a potential synapomorphy supporting their relationship. However, since persistent sepals also occur in some other taxa within Ranunculeae, such as *Beckwithia* Jeps., *Ranunculus glacialis* L., and *R. similis* Hemsl., and dioecious reproduction also occur in *Hamadryas* Comm. ex Juss.of trib. Ranunculeae, the systematic position of *Paroxygraphis* requires further investigation to confirm. Despite extensive molecular phylogenetic studies of trib. Ranunculeae (Johansson 1998; Hörandl et al. 2005; Paun et al. 2005; Lehnebach et al. 2007; Hoot et al. 2008; Gehrke and Linder 2009; Hoffmann et al. 2010; Emadzade et al. 2010; Emadzade and Hörandl 2011; Wang et al. 2014), none have included *Paroxygraphis*, leaving its systematic position unresolved.

**Figure 1.**
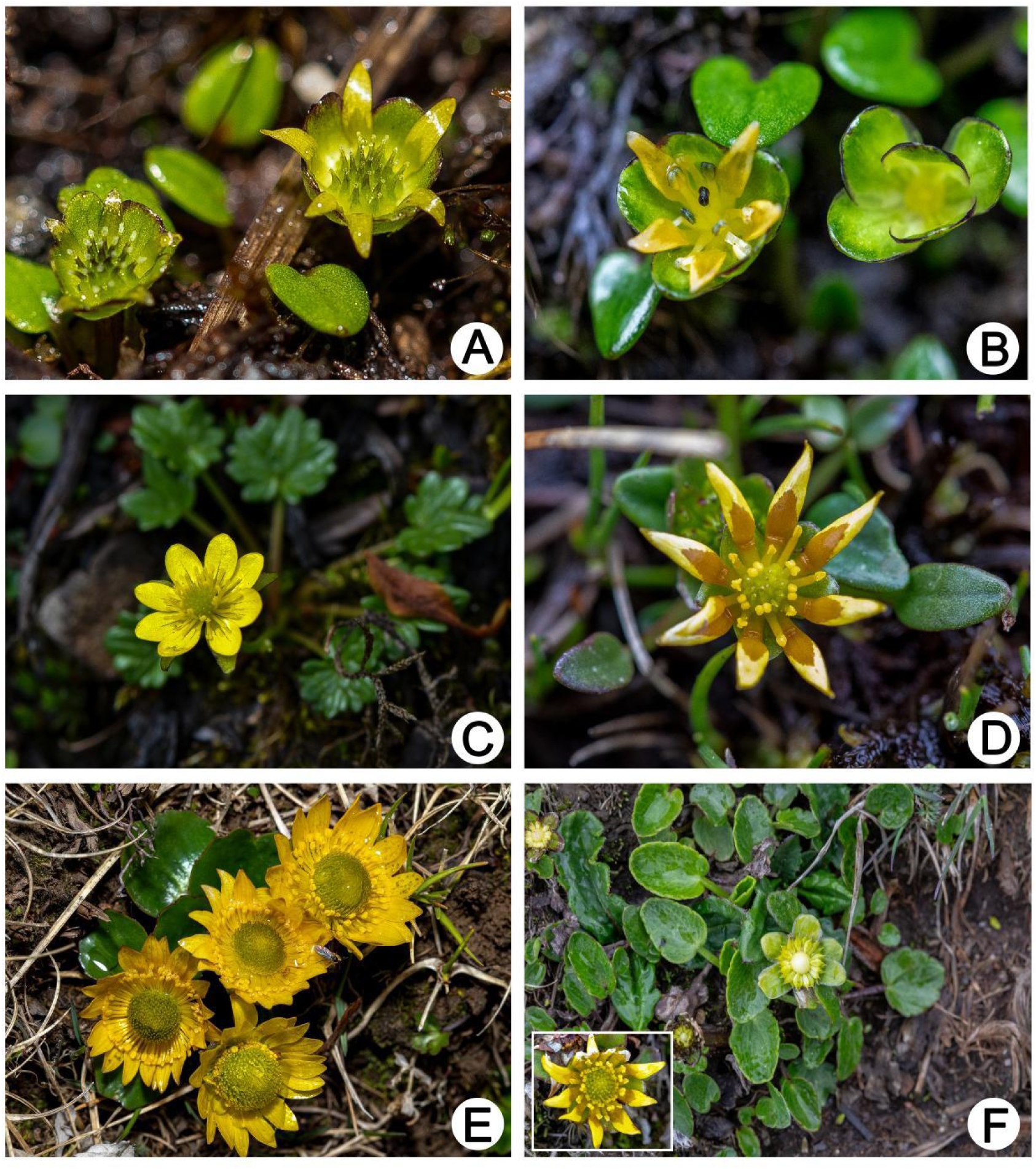
Field photographs of Oxygraphis and Paroxygraphis plants. A–B: Paroxygraphis sikkimensis individuals (photoed by Bo-Wen Liu, in Dinggyê county, Xizang province); (A) Female individual showing pistillate flowers; (B) Male individual with staminate flowers; C: Oxygraphis delavayi in flowering stage (photoed by Lei Xie, in Bomê county, Xizang province); D: Oxygraphis tenuifolia in flowering stage (photoed by Bo-Wen Liu, in Mount Sejila, Nyingchi county, Xizang province); E: Oxygraphis glacialis exhibiting typical morphology (photoed by Bo-Wen Liu, in Mount Mila Pass, Maizhokunggar county, Xizang province); F: Oxygraphis endlicheri displaying flower, fruits, and leaves (photoed by Bo-Wen Liu, in Latola Pass, Gyirong county, Xizang province).

During our spring 2025 field expedition in southern Xizang province, we rediscovered *Paroxygraphis sikkimensis* in Dinggyê county. This finding marks both the first documented rediscovery of the species in recent decades and its first confirmed occurrence within China. The plants were found growing in moss cushions on rocky cliffs at approximately 3800 m elevation, indicating a strict ecological specialization to humid alpine micro-habitats. Specimens were carefully collected for morphological examination and molecular phylogenetic analysis (silica-gel-preserved leaf tissue). This study seeks to: (1) resolve the phylogenetic placement of *Paroxygraphis* within trib. Ranunculeae using molecular systematics, (2) reconstruct its evolutionary history, and (3) reassess its taxonomic classification in light of phylogenetic evidence.

## 2 Materials and Methods

### 2.1 Plant Sampling and Choosing Molecular Markers

In this study, we conducted field surveys and collected leaf samples of *Paroxygraphis sikkimensis* in Dinggyê county, Xizang province, China. Both male and female individuals were separately collected from field. To determine the phylogenetic position of *Paroxygraphis*, we also conducted extensive field surveys and sampling across the genus *Oxygraphis*, obtaining molecular materials from other four recognized species in China for comparative analyses. Species name and delimitation of *Oxygraphis* in this study follow the treatment Flora of China (Wang et al., 2001). It is noteworthy that because the combination of *O. polypetala* (Royle) Hook. f. & Thoms. was based on an illegal basionyme *Ranunculus polypetalus* Royle, this species name has been changed as *O. endlicheri* (Walp.) Bennet & Sum. Chandra (Wang et al. 2001). The recently published new species from India, *O. kumaonensis* I. D. Rai and G. S. Rawat, is not available in this study (Rai and Rawat 2015).

*Ranunculus similis*, which similarly possesses persistent calyx and was previously classified in the genus *Oxygraphis*, was also collected in the field. Besides the above eight samples (six species), we included species from all remaining Ranunculeae genera as ingroups in our molecular phylogenetic analysis. These Ranunculeae sample data were sourced from earlier studies (Wang et al. 2014).

Multiple molecular phylogenetic and phylogenomic studies (Ro et al. 1997; Wang et al. 2009, 2016; Cossard et al. 2016; He et al. 2022) have shown that the trib. Anemoneae is the sister group to the trib. Ranunculeae. Therefore, this study selected species from the genera *Clematis* L., *Anemone* L., *Pulsatilla* Mill., and *Hepatica* Adans. as outgroups. In total, 83 species (representing all 18 genera) and 87 samples of trib. Ranunculeae and four samples of trib. Anemoneae are included in this study (Table S1). Voucher specimens of the newly generated molecular data were deposited in the Herbarium of Beijing Forestry University (BJFC, herbarium code follows Thiers 2017).

We selected a combination of molecular markers including the nuclear ribosomal ITS region and four plastid regions (*mat*K, *psb*J-*pet*A, *rbc*L, and *trn*L-F) according to previously published studies (Wang et al. 2014) to cover all the genera of trib. Ranunculeae. For newly collected samples, we obtained genome skimming data through next-generation sequencing method. Using the aforementioned DNA regions from GenBank as references, we assembled these five target DNA regions from the genome skimming data.

### 2.2 Genome Skimming Sequencing

Total genomic DNA was isolated from silica-dried leaf material at Berry Genomics Co., Ltd. (https://www.bioon.com.cn/company/index/df7a2b444074) using a commercial DNA extraction kit (Tiangen Biotech Co. Ltd., Beijing, China) following the manufacturer’s protocol. For library preparation, 1 μg of DNA per sample was processed with the VAHTS Universal DNA Library Prep Kit for MGI (Vazyme, Nanjing, China) to generate sequencing-ready libraries. Paired-end libraries of 2×150 bp were constructed and sequenced using on a DNBSEQ-T7 platform (BGI, Shenzhen, China). Each sequencing run produced approximately 6 Gbp of raw data per sample.

### 2.3 Phylogenetic Analyses

After assembling the five DNA regions from the newly sequenced species, sequences of each region were aligned using MAFFT v.6.833 (Katoh et al. 2005) with iterative manual optimization in Geneious v.Prime (Kearse et al. 2012), followed by pruning of ambiguous alignment regions according to Wang et al. (2014).

Phylogenetic analyses for nrITS, plastid region, and combined data (congruent test between nrITS and plastid regions followed Wang et al. 2014) were carried out using both maximum likelihood (ML) and Bayesian inference (BI) methods in RAxML v.8.1.17 (Stamatakis 2014) and MrBayes v3.2.3 (Ronquist and Huelsenbeck 2003), respectively. The ML analysis was carried out using a GTR+G substitution model substitution model as recommended in the software documentation. Branch support values in the maximum likelihood phylogeny were assessed through 500 nonparametric bootstrap replicates (MLBS). For BI analyses, the best-fit model of nucleotide substitution, as determined by the Akaike information criterion in jModelTest 2 (Darriba et al. 2012). Four parallel MCMC chains were implemented in our Bayesian inference, beginning with random tree topologies. After running for 2,000,000 generations and sampling every 1000th generation, we excluded the first 20% of trees as burn-in before constructing a >50% majority-rule consensus tree to obtain posterior probability values (PP).

### 2.4 Molecular Dating

Due to the limited phylogenetic information provided by nrITS and plastid data, we used the combined data for molecular dating analysis according to Wang et al. (2014). Our molecular clock analysis incorporated four temporal constraints established by Emadzade and Hörandl (2011): (1) the Ranunculeae-Anemoneae divergence (46.6 Ma; Anderson et al. 2005, molecular estimate); (2) the minimum age constraint for *Myosurus* (23 Ma; Oligocene fossil evidence from Mai and Walter 1978); (3) the *R. carpaticola*-*R. notabilis* speciation event (0.914 Ma; allozyme data from Hörandl 2004); and (4) the maximum age limit for *R. caprarum* diversification (2 Ma; Stuessy et al. 2006). Calibration priors were implemented according to Emadzade and Hörandl’s (2011) specifications.

Divergence times were estimated using a Bayesian approach implemented in BEAST v.2.6 (Bouckaert et al. 2014). The analysis was performed using the GTR+G substitution model, an uncorrelated log-normal relaxed molecular clock, and a Yule pure-birth model as the tree prior. The Markov chain Monte Carlo (MCMC) analyses were run for 200 million generations, with parameters sampled every 10,000 generations. Convergence of the three independent runs was assessed in Tracer v.1.7 (Rambaut et al. 2018) by ensuring all effective sample size (ESS) values were above 200. After confirming convergence, the initial 25% of samples from each run were discarded as burn-in, and the results from the runs were combined.

## 3 Results

### 3.1 Phylogeny of trib. Ranunculeae

The aligned matrix of the concatenated plastid regions consisted of 4664 characters with 1559 variable and 1043 parsimony-informative sites. The aligned matrix of ITS sequences was 599 nucleotides in lengthwith 343 variable and 261 parsimony-informative sites. The ML and BI analyses resulted in identical trees for each data set. Contrary to the plastid topologies, the ITS topology has very low resolution, but the nodes in the ITS tree with PP>0.95 are also recognized by plastid data (Figures S1–S4). The combined matrix included 5263 aligned characters with 1902 variable and 1304 parsimony-informative sites. Since the combined dataset yielded the phylogeny (Figure 2) with both highest generic-level resolution and strongest nodal support, discussions of intergeneric relationships in this study are predominantly based on the concatenated phylogenetic reconstruction.

**Figure 2.**
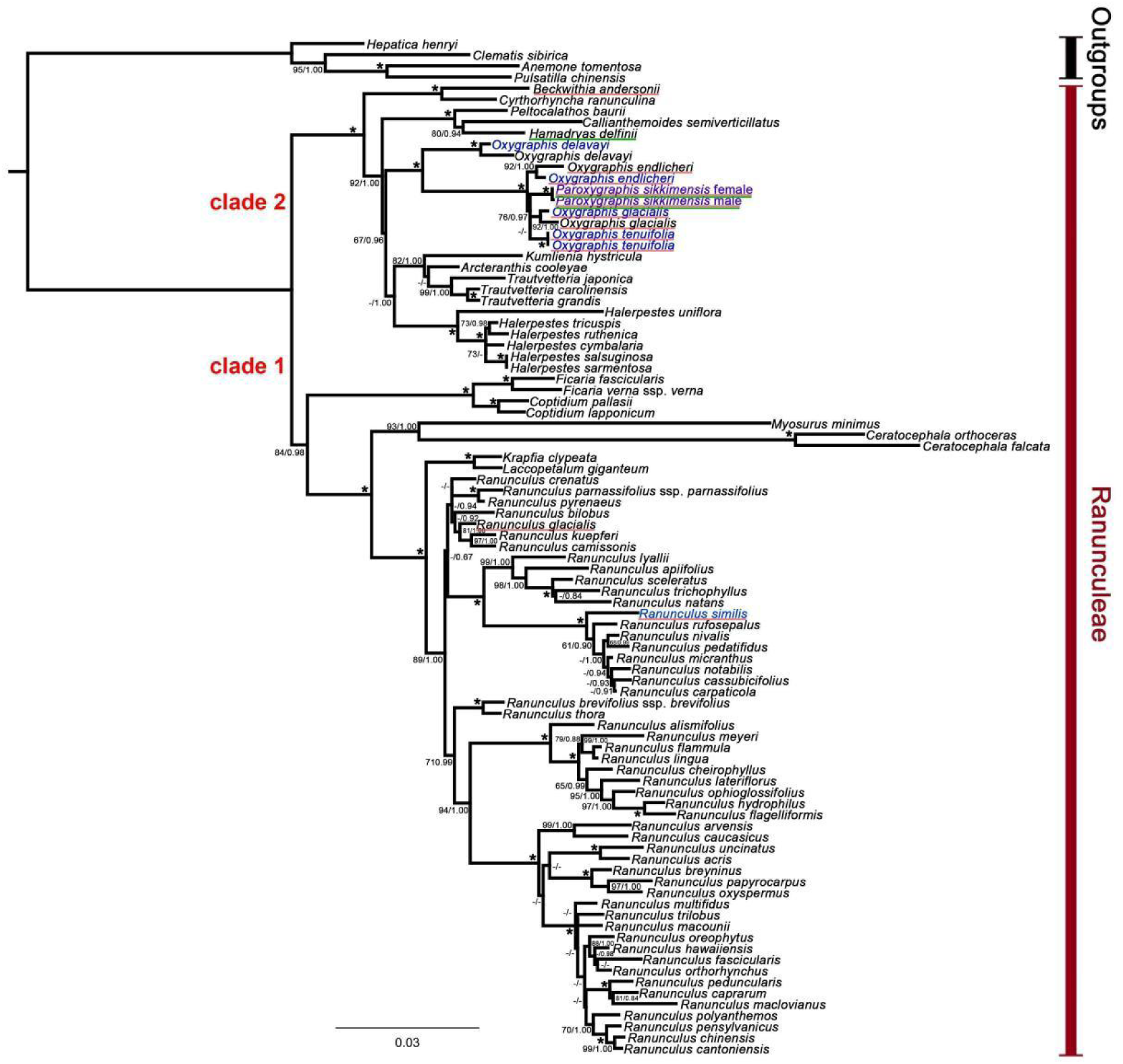
Phylogram of trib. Ranunculeae obtained from the combined datasets. Numbers above and below branches are ML bootstrap (BS) percentages and Bayesian posterior probabilities, respectively. Only nodes with maximum likelihood BS>60, and posterior probability > 0.80 were indicated. Plant names in blue are newly sequenced in this study, and those in black are retrieved from GenBank. Red underlined species are with persistent sepals and green underlined ones are dioecious.

When using Anemoneae species as the outgroup, all 18 sampled genera of trib. Ranunculeae form a strongly supported monophyletic clade (nrITS data MLBS=100, PP = 1.0; plastid data: MLBS=100, PP = 1.0; combined data: MLBS=100, PP = 1.0, Figures 2, S1–S4). Our phylogenetic analyses (the plastid and the combined datasets, but not nrITS) resolved two major clades within trib. Ranunculeae (Figures 2, S1–S2). Clade I comprised *Ranunculus* along with *Laccopetalum* Ulbr., *Krapfia* DC., *Ceratocephala* Moench, *Myosurus*, *Coptidium* (Prantl) Beurl. ex Rydb., and *Ficaria* Schaeff., while Clade II included *Trautvetteria* Fisch. et C.A. Mey., *Arcteranthis* Greene, *Kumlienia* Greene, *Halerpestes* Greene, *Oxygraphis* (containing *Paroxygraphis*), *Callianthemoides* Tamura, *Hamadryas* Tamura, *Peltocalathos* Tamura, *Cyrtorhyncha* Nutt., and *Beckwithia*. In the chloroplast gene tree (Figures S1–S2), the *Oxygraphis* clade formed a sister-group relationship with *Arcteranthis*, albeit with weak support (BS=63, PP = 1.0). Combined data (Figure 2) revealed that the *Oxygraphis* clade forms a sister relationship with a branch comprising *Kumlienia*, *Arcteranthis*, *Halerpestes*, and *Trautvetteria* (BS=67%, PP=0.96).

The nrITS sequences failed to resolve the *Oxygraphis* clade (Figures S3–S4). *Oxygraphis delavayi* Franch. (the only *Oxygraphis* species without persistent sepals) was separated from the other species of *Oxygraphis*, but with low overall support. However, *Paroxygraphis* formed a strongly supported clade with all the other *Oxygraphis* species in nrITS tree (MLBS=100, PP=1). In this clade, all the species have persistent sepals.

The plastid and combined datasets consistently place both male and female individuals of *P. sikkimensis* within the *Oxygraphis* clade (Figures 2, S1–S2, combined data MLBS=100, PP=1; plastid data MLBS=100, PP=0.98). Within this clade, *O. delavayi* represents the earliest-diverging lineage and is uniquely characterized by its lack of persistent sepals, a morphological character retained in all other congeneric species as well as *P. sikkimensis*. Within the clade of persistent-sepal *Oxygraphis* species (Figure 2, combined data MLBS=100, PP=1), *O. endlicheri* diverged first, while *O. tenuifolia* W. E. Evans, *O. glacialis* Regel, and *Paroxygraphis* formed a subclade with unresolved internal relationships.

Within Ranunculeae, the other dioecious genus *Hamadryas* forms a sister-group relationship with *Callianthemoides* (BS=80, PP=0.94, Figure 2), while showing no direct ancestral connection with *Paroxygraphis*. *Beckwithia*, another genus characterized by persistent sepals, forms a sister-group relationship with *Cyrtorhyncha* (BS=100%, PP=1.0, Figure 2), also showing no direct connection to *Paroxygraphis*.

### 3.2 Molecular Dating

The analyses based on combined dataset resulted in very similar divergence-time estimates (Figure 3, Table 1) for genera of Ranunculeae with previous studies (Wang et al. 2014). Our time estimates suggest that extant Ranunculeae date to Eocene (node 1, 46.72 Ma) (95% highest posterior density, HPD: 44.76–48.65 Ma). The generic diversification of clade I occurred in the Late Eocene (node 9, 38.92 Ma, 95% HPD: 32.58–44.97 Ma). The largest genus in Ranunculeae, *Ranunculus*, diversified (crown age) at 23.88 Ma (node 11, 95% HPD: 17.56–30.11 Ma). The age of crown clade II is estimated to be 24.95 Ma (node 3, 95% HPD: 17.21–33.17 Ma). The *Oxygraphis* clade diverged from its close relatives at 19.54 Ma (node 5, 95% HPD: 13.61–25.78 Ma), and the current species of this clade diversified at 13.75 Ma (95% HPD: 8.26–19.70 Ma). In the *Oxygraphis* clade, the crown group age of the subclade containing species with persistent sepals is 4.53 Ma (node 7, 95% HPD: 2.51–6.86 Ma). *Paroxygraphis sikkimensis* split from its sister clade at 3.73 Ma (95% HPD: 1.98–5.77 Ma).

**Figure 3.**
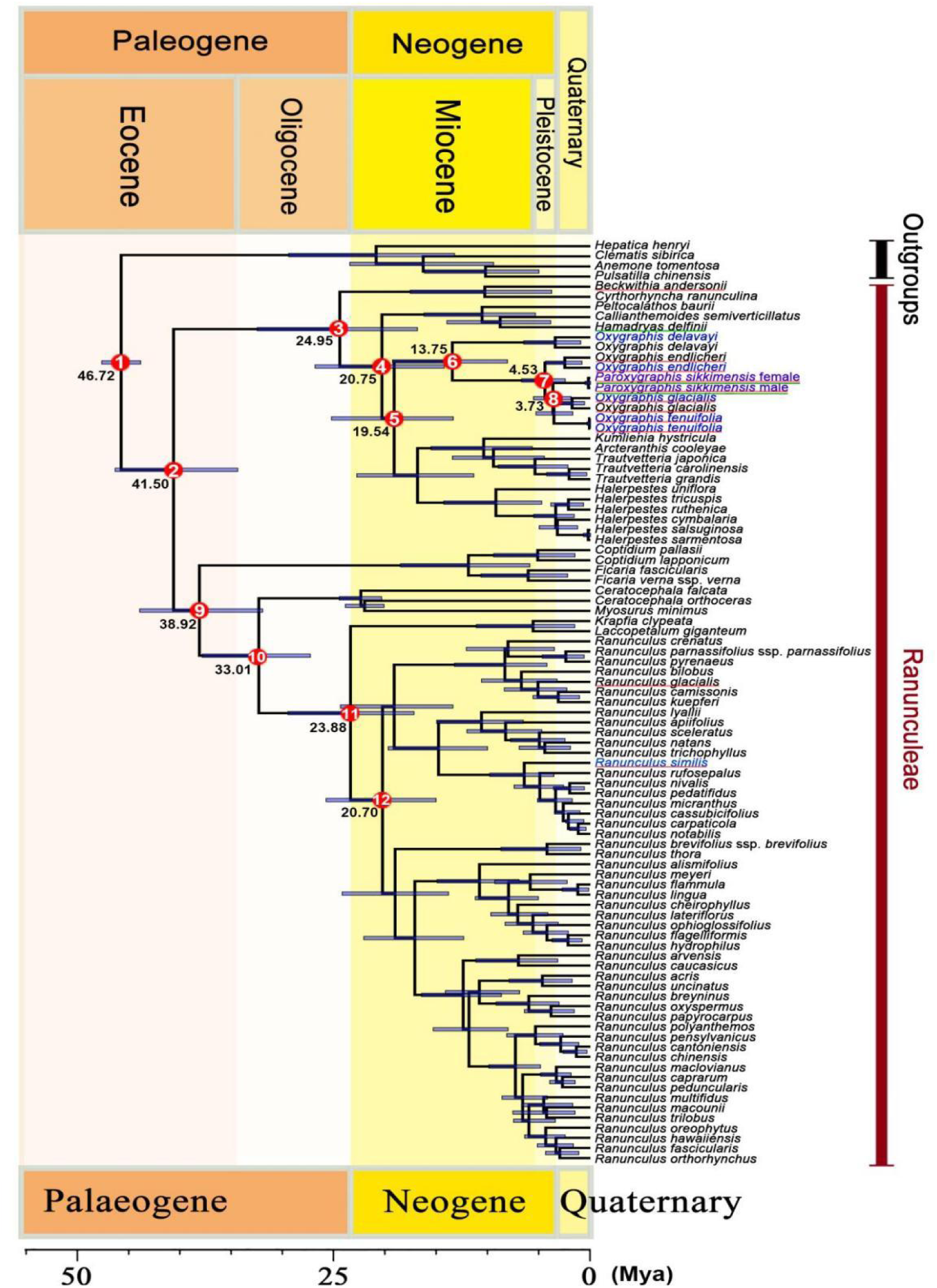
Maximum clade credibility chronogram of trib. Ranunculeae using combined datasets. Numbered red circles indicate nodes for which full divergence time statistics are detailed in Table 1; the adjacent number represents the mean age estimate (Ma) for that node. Plant names in blue are newly sequenced in this study, and those in black are retrieved from GenBank. Red underlined species are with persistent sepals and green underlined ones are dioecious.The geological time scale (Gradstein et al., 2004) is shown at the bottom.

**Table 1.** Divergence time estimates (Ma) for trib. Ranunculeae. The node numbers are shown in Figure 3.

| Nodes | Mean Height (Ma) | Median Height (Ma) | Lower 95% HPD (Ma) | Upper 95% HPD (Ma) | Posterior Probability |
| --- | --- | --- | --- | --- | --- |
| 1 | 46.72 | 46.72 | 44.76 | 48.65 | 1.00 |
| 2 | 41.5 | 41.82 | 35.11 | 47.31 | 1.00 |
| 3 | 24.95 | 24.63 | 17.21 | 33.17 | 1.00 |
| 4 | 20.75 | 20.51 | 14.55 | 27.4 | 1.00 |
| 5 | 19.54 | 19.32 | 13.61 | 25.78 | 0.96 |
| 6 | 13.75 | 13.52 | 8.26 | 19.70 | 1.00 |
| 7 | 4.53 | 4.39 | 2.51 | 6.86 | 1.00 |
| 8 | 3.73 | 3.61 | 1.98 | 5.77 | 0.97 |
| 9 | 38.92 | 39.02 | 32.58 | 44.97 | 0.96 |
| 10 | 33.01 | 32.85 | 27.88 | 38.64 | 1.00 |
| 11 | 23.88 | 23.76 | 17.56 | 30.11 | 1.00 |
| 12 | 20.7 | 20.56 | 15.37 | 26.31 | 1.00 |

## 4 Discussion

### 4.1 Phylogenetic Relationships Within trib. Ranunculeae

Our plastid and combined datasets yielded phylogenetic relationships that are largely congruent with and of comparable reliability to those reported in previous studies (Emadzade et al. 2010, 2011; Emadzade and Hörandl 2011; Wang et al. 2014). The well-supported clades we recovered align closely with established topologies. However, due to the inherent limitations of Sanger sequencing data in providing sufficient phylogenetic signal, the systematic positions of certain genera, including *Arcteranthis*, *Hamadryas*, and *Peltocalathos*, remain unresolved and warrant further investigation.

Consistent with prior work, our analyses resolved tribe Ranunculeae into two major clades (Emadzade et al. 2010; Wang et al. 2014). While phylogenetic relationships among genera within Clade I were relatively well-resolved, those within Clade II exhibited lower resolution, reflecting deeper uncertainties in intergeneric relationships (Figures 2–3). Notably, our results place the *Oxygraphis* clade within Clade II, yet its closest relatives remain ambiguous. The combined dataset weakly suggests a sister relationship between the *Oxygraphis* clade and a clade comprising *Kumlienia*, *Arcteranthis*, *Trautvetteria*, and *Halerpestes*, whereas plastid data alone tentatively support *Arcteranthis* as sister to *Oxygraphis* (BS = 56, PP = 0.98).

Our findings in Clade II neither fully contradict nor entirely align with previous studies (Emadzade et al. 2010, 2011; Emadzade and Hörandl 2011; Wang et al. 2014). These discrepancies likely arise from the limited phylogenetic informativeness of Sanger sequencing data, which proved insufficient for robustly resolving recent divergences within Clade II. In contrast, the deeper divergence times among genera in Clade I facilitated clearer phylogenetic inference. Given these limitations, the precise placement of *Oxygraphis* will require higher-resolution genomic-scale data in future studies.

### 4.2 Origin and Evolution of *Oxygraphis* and *Paroxygraphis*

*Oxygraphis* is a genus within clade II of trib. Ranunculeae that exhibits special adaptations to alpine and arctic environments. All four or five species in this genus can be found in the Hengduan Mountains, the Qinghai-Xizang Plateau, and Himalayan regions, with three or four of them being endemic to this region (Wang et al. 2001; Rai and Rawat 2015). Only *O. glacialis* extends further into the cold regions of northern Asia, with its range reaching as far as Alaska in North America. Although polyploidization is prevalent in many taxa within trib. Ranunculeae (Hörandl et al. 2005; Paun et al. 2005, 2006; Baltisberger and Widmer 2009; Cires et al. 2013), all currently reported *Oxygraphis* species are diploid (2n=16, x=8, Yang 2000).

As observed in prior studies (Wang et al., 2014), *O. delavayi* (with deciduous sepals) is the earliest diverging lineage within the genus (Figures 2–3, Table 1), and its divergence time from the species clade with persistent-sepal is relatively old in Miocene, reaching 13.75 Ma (95% HPD: 8.26–19.70 Ma). The persistent-sepal clade diversified later, during the Pliocene (4.53 Ma, 95% HPD: 2.51–6.86 Ma), coinciding with severe cooling in Asia. This climatic shift may have driven the evolution of persistent sepals in *Oxygraphis* to safeguard their fruits.

The results of this study indicate that *Paroxygraphis* is embedded within the *Oxygraphis* clade, with an estimated divergence time from its closest sister group of approximately 3.73 Ma (95% HPD: 1.98–5.77 Ma). *Paroxygraphis sikkimensis* is aslo an alpine and cold-adapted species, restricted to high-elevation zones of the southeastern Himalayas. Unlike most *Oxygraphis* species inhabiting alpine meadows, *P. sikkimensis* grows exclusively on moss-covered, damp rock crevices at elevations of about 3,800 m. Due to its diminutive size, early anthesis (May), and special habitat, this species is exceptionally hard to find and easily overlooked in field surveys.

*Paroxygraphis sikkimensis* typically bears only 5–6 petals, significantly fewer than other species in the genus *Oxygraphis* (Tamura 1995, Figure 1). However, its persistent sepals represent a shared characteristic with three or four *Oxygraphis* species. Our phylogenetic analyses (even in nrITS phylogeny) reveal that this persistent sepal trait constitutes a synapomorphy for four species (*O. glacialis*, *O. tenuifolia*, *O. endlicheri*, and *P. sikkimensis*) within the *Oxygraphis* clade, excluding only the earliest diverging member, *O. delavayi*. However, within trib. Ranunculeae, the trait of persistent sepal has evolved independently in multiple lineages (Figures 2–3). This characteristic is also present in *Beckwithia*, *Ranunculus glacialis*, and *Ranunculus similis*. Emadzade et al. (2010) noted that these taxa all occur in high alpine or Arctic zones, suggesting their persistent sepals may represent a homoplastic adaptation protecting developing fruits from wind and extreme cold in harsh environments.

*Paroxygraphis* is morphologically highly similar to *Oxygraphis*, but traditional classifications consistently treated it as a distinct genus primarily based on its dioecious reproductive system (Smith 1913; Tamura 1995; Emadzade et al. 2010). Within trib. Ranunculeae, the South American genus *Hamadryas* similarly exhibits a dioecious sexual system (Tamura 1995; Hoot et al. 2008). Our molecular phylogenetic results demonstrate that *Hamadryas* and *Callianthemoides* (Figures 2–3) form a sister group, sharing no direct common ancestry with *Paroxygraphis*. This indicates that dioecy has evolved independently at least two times within trib. Ranunculeae. Moreover, certain genera within Ranunculaceae, such as *Clematis* L. and *Thalictrum* L., contain both hermaphroditic species and dioecious species, demonstrating sexual system variability within these genera. Considering the robust molecular phylogenetic relationships revealed by our analyses, we recommend recognizing *Paroxygraphis* as congeneric with *Oxygraphis*.

At the time of its publication (Smith 1913), *Paroxygraphis sikkimensis* was described with reference to multiple herbarium specimen gatherings but without explicit designation of a holotype. These specimens collectively constitute syntypes under Articles 9.5 and 40.2 of the International Code of Nomenclature (Turland et al. 2018). Herein, we formally designate specimen *Smith 3204* (K000692711) at the Kew Herbarium as the lectotype to stabilize the taxonomic application of this name. The selected lectotype best represents the species’ diagnostic characters as described in the protologue and shows optimal preservation of key morphological features.

## 5 Taxonomic treatment

**Oxygraphis sikkimensis** (W. W. Smith) Wen He Li & L. Xie, **comb. nov.**

**Basionym**: *Paroxygraphis sikkimensis* W. W. Smith, Rec. Bot. Surv. India 4: 344. 1913.

**Type**: India, Sikkim, near Changu bungalow, in the Dikchu valley, in the Chola Range below the Tanka La, in the wetter ranges at 12000-14000 ft, *Smith 3204* [**lectotype here desinated**: K (K000692711!); isolectotype: E!]

**Additional specimen cited:** China, Xizang Province, Dinggyê County, near the China-Nepal border (Due to the species’ extremely limited distribution and conservation concerns, precise geographic coordinates have been intentionally omitted from this publication to prevent anthropogenic disturbances.), alt. 3850m, 18 May, 2025, *Wen-He Li & Bo-Wen Liu 2025051806* (BJFC); alt 3800m, 19 May, 2025, *Wen-He Li & Bo-Wen Liu 2025051905* (BJFC).

**Ecology, distribution and status:** This species was known only from a few localities in India (Sikkim) and Nepal (Smith 1913), and is now reported in southern Xizang province of China. In Dinggyê population, it grows in moist moss cushions on wet cliffs. The area we discovered is rather small, including a few hundred individuals.

This species’ habitat is highly vulnerable to anthropogenic disturbances, which may lead to population degradation or even local extinction. According to the IUCN red list categories and criteria (IUCN Standard and Petitions Committee 2022), *Oxygraphis sikkimensis* should be categorized as critically endangered (CR).

## 6 Conclusions

Our integrative study resolves the century-old taxonomic ambiguity surrounding *Paroxygraphis* by demonstrating its nested position within *Oxygraphis*, supported by molecular phylogenetics, morphological synapomorphies, and biogeographic evidence. The rediscovery of *O. sikkimensis* in Xizang expands its known range and underscores the Himalayan region as a hotspot for alpine plant diversification.

Convergent evolution of persistent sepals and dioecy in trib. Ranunculeae highlights adaptive responses to harsh environments. Taxonomic reclassification of *Paroxygraphis* under *Oxygraphis* rationalizes its phylogenetic affinity while preserving diagnostic traits. However, the species’ extreme habitat specialization and limited distribution necessitate urgent conservation measures. This work not only advances systematic understanding of trib. Ranunculeae but also emphasizes the importance of combining field exploration with molecular tools to unravel cryptic biodiversity in alpine ecosystems. The discovery of *Oxygraphis sikkimensis* further suggests that our understanding of the flora in China’s Himalayan regions remains incomplete.

## Author Contributions

Wen-He Li: data curation (equal), formal analysis (equal), field investigation (lead), methodology (equal), resources (lead), software (equal), validation (lead), visualization (lead), writing – original draft (equal), writing – review and editing (equal). Wan-Ting Chen: data curation (Lead), resources (lead), writing – review and editing (equal). Bo-Wen Liu: data curation (equal), methodology (equal), field investigation (lead), software (supporting), visualization (supporting), writing – original draft (supporting), writing – review and editing (supporting). Jia-Min Xiao: data curation (equal), investigation (equal), resources (equal), supervision (supporting), visualization (equal). Huan-Yu Wu: data curation (supporting), investigation (supporting), resources (supporting), supervision (supporting), visualization (equal). Jian He: conceptualization (lead), funding acquisition (equal), project administration (lead), supervision (lead), writing – review and editing (lead). Lei Xie: conceptualization (equal), funding acquisition (lead), project administration (lead), supervision (lead), writing – review and editing (equal).

## Acknowledgments

This research was funded by the Discipline Crossing Foundation of School of Ecology and Nature Conservation, Beijing Forestry University (grant number: BH2025-JX-03); the National Natural Science Foundation of China (grant number: 31670207). We thank Dr. Liang-Liang Yue and Zong-Zong Yang for helping to obtain some *Oxygraphis* samples.

## Data Availability Statement

The DNA sequences generated in the present study have been deposited in the National Center for Biotechnology Information (NCBI) database. The accession numbers and the information on the voucher specimens are available in Table S1. The voucher specimens were housed in BJFC.

## Supplementary material

**Figure S1.** Maximum likelihood (ML) phylogram of tribe Ranunculeae, inferred using RAxML from the plastid dataset. Numbers above and below branches represent ML bootstrap percentages (BS). Support values are shown only for nodes with BS > 60. Taxon names in blue indicate species newly sequenced for this study, while those in black are from sequences retrieved from GenBank. Species with persistent sepals are underlined in red, and dioecious species are underlined in green.

**Figure S2.** Bayesian inference (BI) phylogram of tribe Ranunculeae, inferred from the plastid dataset using MrBayes. Numbers at the nodes represent Bayesian posterior probabilities (PP). Only support values with PP > 0.80 are shown. Taxon names in blue indicate species newly sequenced for this study, while those in black are from sequences retrieved from GenBank. Species with persistent sepals are underlined in red, and dioecious species are underlined in green.

**Figure S3.** Maximum likelihood (ML) phylogram of tribe Ranunculeae, inferred using RAxML from the ITS dataset. Numbers above and below branches represent ML bootstrap percentages (BS). Support values are shown only for nodes with BS > 60. Taxon names in blue indicate species newly sequenced for this study, while those in black are from sequences retrieved from GenBank. Species with persistent sepals are underlined in red, and dioecious species are underlined in green.

**Figure S4.** Bayesian inference (BI) phylogram of tribe Ranunculeae, inferred from the ITS dataset using MrBayes. Numbers at the nodes represent Bayesian posterior probabilities (PP). Only support values with PP > 0.80 are shown. Taxon names in blue indicate species newly sequenced for this study, while those in black are from sequences retrieved from GenBank. Species with persistent sepals are underlined in red, and dioecious species are underlined in green.

**Table S1** Species, GenBank accession numbers, vouchers, of the sequences used in this study.

