## Supplementary figures and images for "Rediscovery of the Enigmatic *Paroxygraphis* in Xizang, China: Phylogenetic Evidence for its Reclassification Within *Oxygraphis* (Ranunculaceae)"

### Figure S1

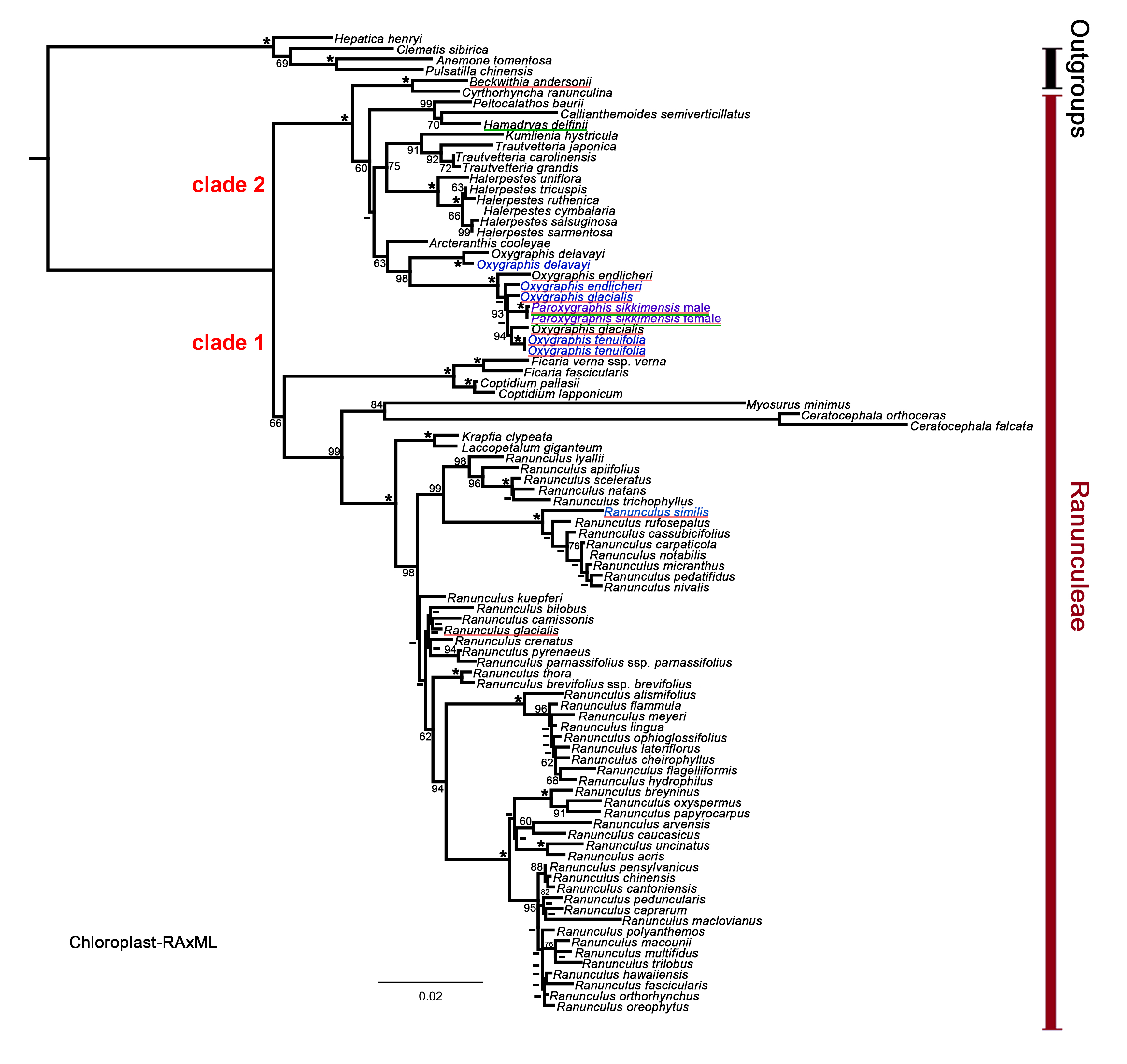

### Figure S2

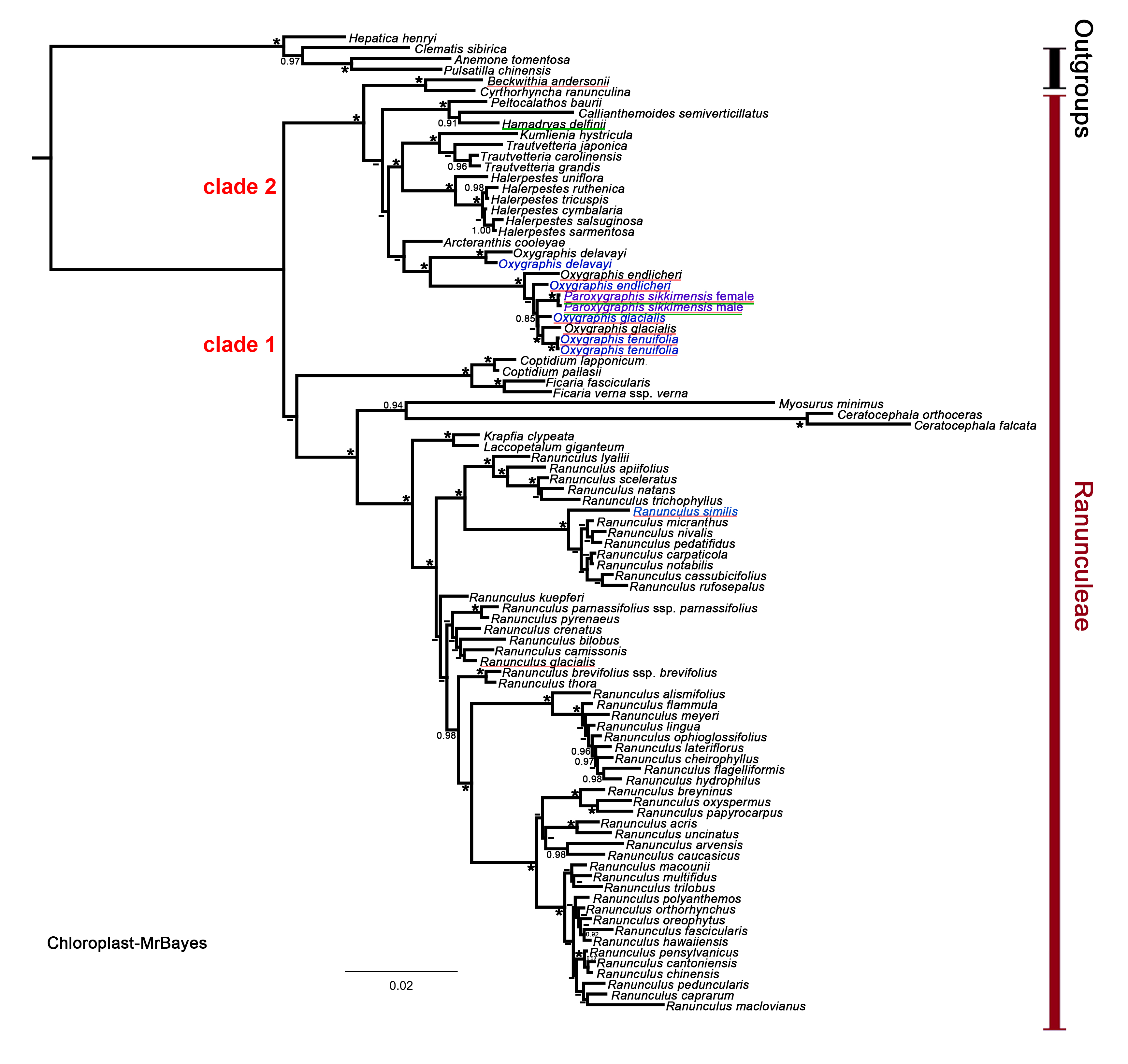

### Figure S3

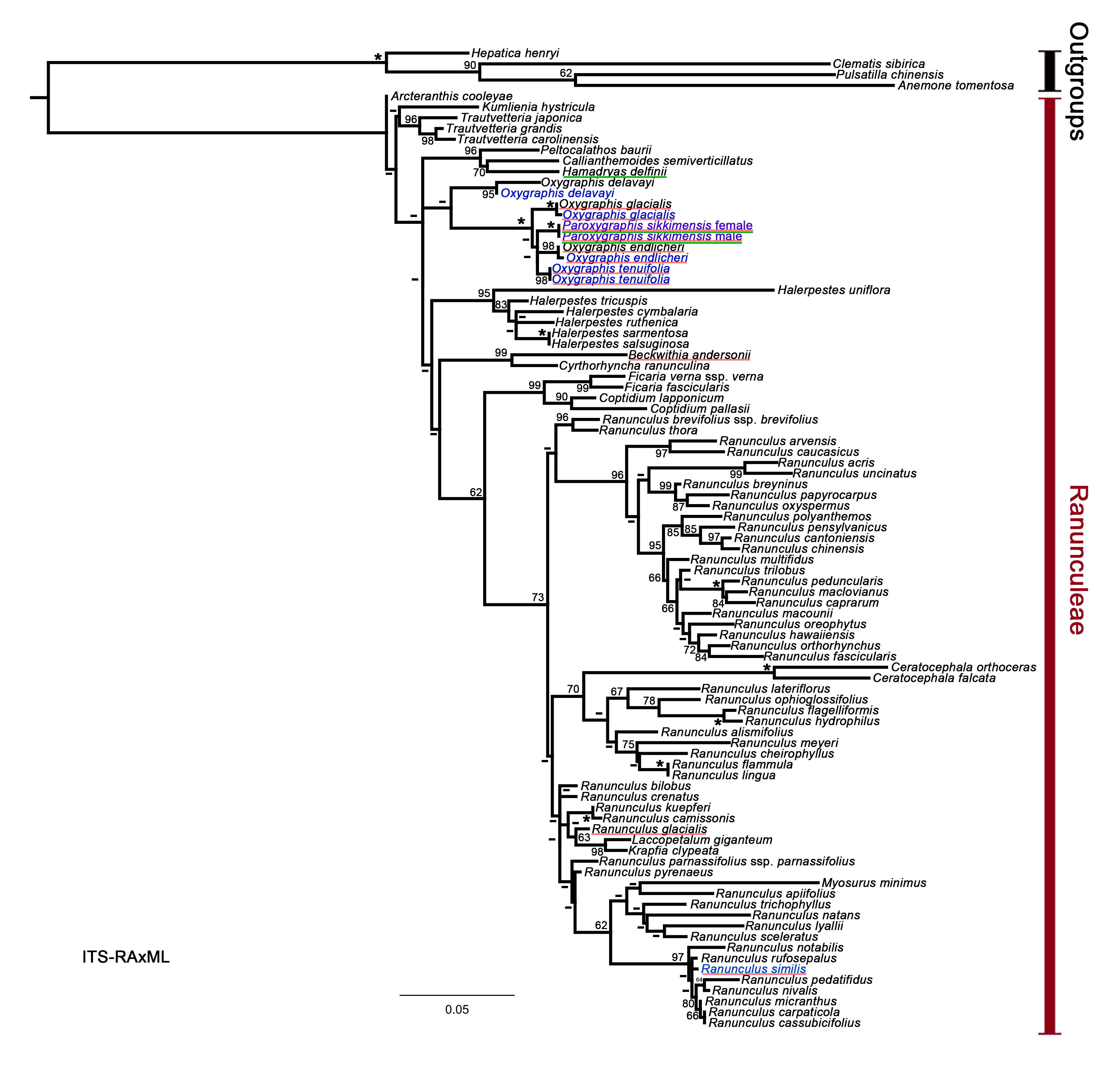

### Figure S4

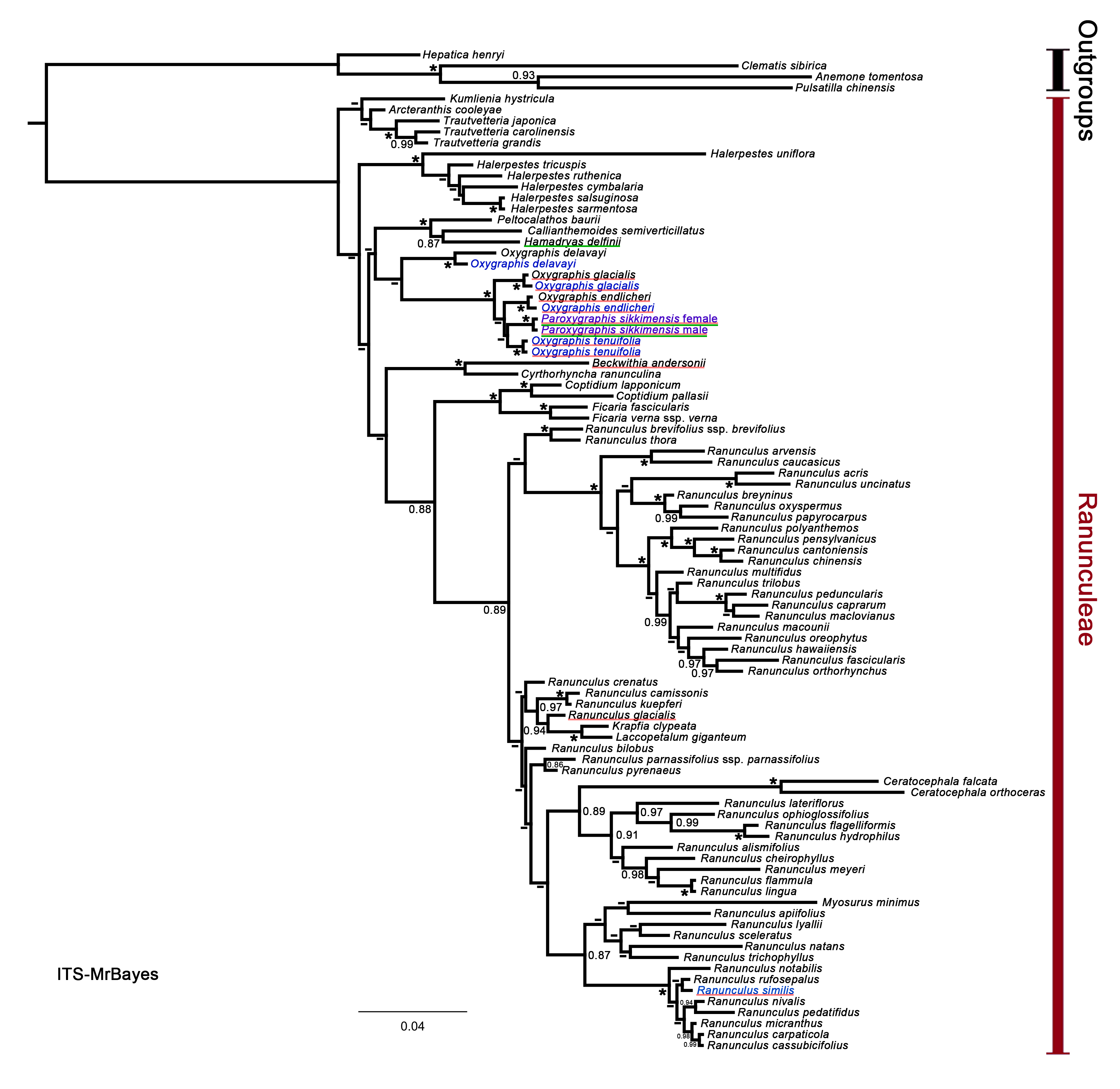
